# Realistic fisheries management reforms could mitigate the impacts of climate change in most countries

**DOI:** 10.1101/804831

**Authors:** Christopher M. Free, Tracey Mangin, Jorge García Molinos, Elena Ojea, Christopher Costello, Steven D. Gaines

## Abstract

Although climate change is altering the productivity and distribution of marine fisheries, climate-adaptive fisheries management could mitigate many of the negative impacts on human society. We forecast global fisheries biomass, catch, and profits to 2100 under three climate scenarios (RCPs 4.5, 6.0, 8.5) and five levels of management reform to (1) determine the impact of climate change on national fisheries and (2) quantify the national-scale benefits of implementing climate-adaptive fisheries reforms. Management reforms accounting for shifting productivity and shifting distributions would yield higher catch and profits in the future relative to today for 60-65% of countries under the two least severe climate scenarios but for only 35% of countries under the most severe scenario. Furthermore, these management reforms would yield higher cumulative catch and profits than business-as-usual management for nearly all countries under the two least severe climate scenarios but would yield lower cumulative catch for 40% of countries under the most severe scenario. Fortunately, perfect fisheries management is not necessary to achieve these benefits: transboundary cooperation with 5-year intervals between adaptive interventions would result in comparable outcomes. However, the ability for realistic management reforms to offset the negative impacts of climate change is bounded by changes in underlying biological productivity. Although realistic reforms could generate higher catch and profits for 23-50% of countries experiencing reductions in productivity, the remaining countries would need to develop, expand, and reform aquaculture and other food production sectors to offset losses in capture fisheries. Still, climate-adaptive management is more profitable than business-as-usual management in all countries and we provide guidance on implementing – and achieving the benefits of – climate-adaptive fisheries reform along a gradient of scientific, management, and enforcement capacities.

## Introduction

Marine fisheries provide a vital source of food for over half the world’s population and support the livelihoods of over 56 million people globally [1]. However, the ability for marine fisheries to provide these services is threatened by climate change [2], compromising the contribution of the oceans to sustainable development goals [3]. Ocean warming has already reduced the productivity of many fisheries around the globe, with some regions having experienced up to 35% declines in maximum sustainable yield [4]. An ensemble of marine ecosystem models forecasts continued decreases in marine animal biomass of 4.8% to 17.2% by 2100 under low- to high-end emissions scenarios, respectively [5]. In general, productivity is predicted to decrease in tropical and temperate regions and increase towards the poles [5], as marine organisms shift distributions to maintain their thermal niches [6–8]. These regional shifts in productivity, range, and fishing opportunity will result in regional discrepancies in food and profits from fisheries [9]. Under current policies, these effects will be unevenly distributed with tropical developing countries and small island developing states exhibiting the greatest vulnerability to the impacts of climate change on fisheries [10–12].

The response of fishers and managers to these changes could either exacerbate or mitigate the impacts of climate change on human society and must be considered in forecasts of climate impacts on marine fisheries [13, 14]. For example, a failure to reduce harvest rates in response to decreasing productivity could increase the risk of overfishing [15], which could subsequently reduce the resilience of stocks to climate change [4] and result in reduced long-term yields [16]. Similarly, a failure to establish transboundary institutions for managing stocks shifting distributions across territorial boundaries could result in the degradation of management and stock health, catch, and profits [17, 18]. In both cases, failing to adapt fisheries management to climate change would exacerbate the impacts of the underlying shifts in productivity on human society. On the other hand, jointly reforming fisheries management and adapting it to account for these climate-driven shifts in productivity and distribution could reduce, or even reverse, the negative impacts of climate change on communities dependent on fishing [13, 19].

Gaines et al. [19] provided a critical step towards understanding the opportunities for fisheries reforms to mitigate the impacts of climate change at a global-level. They showed, at a global scale, that business-as-usual fisheries management would exacerbate the negative impacts of climate change, but that climate-adaptive fisheries reforms would maintain global fisheries health, harvest, and profits into the future under all but the most severe emissions scenario evaluated (RCP 8.5). However, the effectiveness and feasibility of these reforms is likely to vary regionally, with higher capacity, poleward countries gaining productivity and lower capacity, tropical countries losing productivity. Furthermore, the benefits documented by Gaines et al. [19] are likely optimistic, as they assume real-time adaptations to shifting productivity. This degree of adaptation potential is unlikely even in the United States where stock assessments are conducted every two to five years [20] and do not frequently include environmental or ecosystem information [21]. Thus, a critical next step in understanding the potential for fisheries reform to mitigate the impacts of climate change on human livelihoods is to examine the performance of more realistic productivity adaptations at the country-level.

Here, we use the Gaines et al. [19] climate-linked bioeconomic model to evaluate the impacts of climate change and management reform on fisheries representing 156 countries, 779 marine fish and invertebrate species, and approximately 58.2% of reported global catch (45.6 of 78.4 mt in 2012; [1]). The evaluated management scenarios address shifting productivity and distributions along a gradient from no adaptation (a.k.a., business-as-usual management) to full adaptation, including scenarios with realistic intervals between management interventions. Overall, we (1) forecast the impacts of climate change on national fisheries and (2) quantify the national-scale benefits of implementing climate-adaptive fisheries reforms. We conclude with a brief overview of promising methods for achieving the benefits of climate-adaptive fisheries reform along a gradient of scientific, management, and enforcement capacities.

## Methods

### Overview

We used the Gaines et al. [19] climate-linked fisheries bioeconomic model to examine country-level changes in fisheries status, catches, and profits under three emissions scenarios (RCPs 4.5, 6.0, and 8.5; Table S1) and five management scenarios (Table 1) from 2012 to 2100. Gaines et al. [19] evaluated the 915 single- and mixed-species stocks from Costello et al. [16] with the data required to assess current status and forecast future distributions. In this analysis, we evaluated only the 779 single-species stocks, because the spatial distributions of the mixed-species stocks could not be projected by Gaines et al. [19] and therefore could not be spatially allocated into country jurisdictions. Projections began in 2012 with initial biomasses, fishing mortalities, and conditions (i.e., B/B_MSY_) determined by aggregating values from Costello et al. [16] and initial distributions determined by AquaMaps [22]. Projections were made through 2100 using the following general procedure: (1) distributions were updated based on a modified version of the García Molinos et al. [23] species distribution model (see below); (2) carrying capacities were assumed to change in proportion to changes in range size, i.e., a 10% increase in range size results in a 10% increase in carrying capacity; and (3) biomass, catch, and profits were then updated based on a modified version of the Costello et al. [16] bioeconomic model and the selected management scenario. We provide brief descriptions of the species distribution and bioeconomic models below, but see Gaines et al. [19] and the original references for more details.

**Table 1.** Fisheries management scenarios evaluated in the analysis (HCR=harvest control rule; EEZ=exclusive economic zone).

| Management scenario |
| --- |
| **Business-as-usual (no adaptation)**<br>This scenario assumes that no action is taken: management fails to account for range or productivity shifts or fix economically sub-optimal harvest rates. Thus, current fishing mortality is maintained for all static (non-shifting) stocks and gradually shifts to open access for all transboundary (shifting) stocks given the lack of transboundary agreements.<br><u>HCR for static stocks:</u> Current fishing mortality<br><u>HCR for transboundary stocks:</u> Gradual shift from current to open access fishing mortality |
| **Range shift adaptation only**<br>This scenario assumes that management adapts to spatial changes in range location by implementing transboundary institutions that facilitate continued management of stocks as they shift into and out of EEZs. However, management does not address corresponding changes in productivity or fix economically sub-optimal harvest rates. Thus, the scenario prevents open access fishing of transboundary (shifting) stocks but does not otherwise improve fisheries management.<br><u>HCR for static and transboundary stocks:</u> Current fishing mortality |
| **Productivity shift adaptation only**<br>This scenario assumes that management is naturally adaptive to changes in productivity and fixes economically sub-optimal harvest rates by adopting an economically optimal HCR where the appropriate harvest rate depends on the available biomass. However, this scenario assumes that management does not address transboundary issues associated with spatial range shifts. Thus, this scenario optimizes harvest for static (non-shifting) stocks but sees a shift from optimal to open access harvest for transboundary (non-shifting) stocks.<br><u>HCR for static stocks:</u> Economically optimal fishing mortality<br><u>HCR for transboundary stocks:</u> Gradual shift from economically optimal to open access fishing mortality |
| **Full adaptation**<br>This scenario assumes that management fixes economically sub-optimal harvest rates accounting for shifts in productivity and effectively prepares for range shifts by implementing transboundary institutions. Thus, this scenario assumes adaptive, economically optimal harvest rates even as stocks shift into and out of EEZs.<br><u>HCR for static and transboundary stocks:</u> Economically optimal fishing mortality |
| **Realistic adaptation (implemented at 5, 10, and 20-year intervals)**<br>This scenario implements a more realistic representation of the full adaptation scenario by acknowledging that management rarely acts annually. Instead, this scenario assumes that management sets an economically optimal harvest rate accounting for the available biomass at regular assessment intervals and maintains this rate, regardless of shifts in productivity, until the next assessment. The scenario assumes that transboundary institutions maintain this management interval as stocks shift into and out of EEZs.<br><u>HCR for static and transboundary stocks:</u> Economically optimal rate in the year of assessment is maintained until the next assessment (5, 10, or 20 years later) |

### Species distribution model

The modified García Molinos et al. [23] species distribution model (SDM) is a bioclimatic envelope model that uses information on species depth preferences, thermal tolerances, and the direction and speed of thermal change, i.e., climate velocity, to project changes in species distributions under warming. AquaMaps species distribution maps [22] were used as the starting point (i.e., 2012) for the projections. In each subsequent time step, the SDM calculated the relocation of the distribution (thermal envelope) of each species as dictated by the spatial direction and rate of change of local (1° resolution) climate velocities based on sea surface temperatures under the selected emissions scenario (RCPs 4.5, 6.0, and 8.5; Table S1). Range projections are restricted by species’ thermal tolerances and depth preferences [19].

### Bioeconomic model and management scenarios

The modified Costello et al. [16] bioeconomic model uses a Pella-Tomlinson [24] surplus production model to forecast fish population dynamics under five management scenarios (Table 1). The Pella-Tomlinson production model requires four input parameters for each stock: the initial biomass, carrying capacity (*K*), intrinsic growth rate (*g*), and a shape parameter (*ϕ*) that determines the proportion of carrying capacity at which production is maximized. Parameters were developed for species-stocks following the procedure detailed in Gaines et al. [19] and are based on individual stock parameters [16] sourced from a combination of production models fit to the RAM Legacy Database [25] and catch-MSY models [26] fit to the FAO Catch Database [1]. The shape parameter is fixed at the meta-analytic average for fish [27], which maximizes productivity at 40% of carrying capacity. Carrying capacity is updated each year based on the resulting changes in range size from the SDM assuming a 1:1 proportional change (see [19] for a justification of this assumption).

The harvest rate is based on the following five management scenarios: business-as-usual (i.e., no adaptation), productivity shift adaptation only, range shift adaptation only, full adaptation, and “realistic” adaptation (see Tables 1 and 2 for details). Productivity shift adaptations improve fisheries management by implementing a dynamic, economically-optimal harvest policy given current biological conditions, which optimally adjusts harvest mortality on the basis of available biomass and is therefore naturally adaptive to climate-driven productivity changes. Range shift adaptations assume that transboundary cooperation results in the maintenance of management, rather than the degradation of management to open access, as stocks shift across boundaries. Business-as-usual management fails to implement either adaptation: it maintains current harvest rates for species that do not shift spatially, while management degrades to open access for stocks that shift across boundaries. Full adaptation assumes that both challenges are addressed: the dynamic economically-optimal harvest policy is implemented and maintained even as stocks shift across boundaries. Realistic adaptation refines the full adaptation scenario by implementing productivity shift adaptations at plausible management intervals: it determines the economically-optimal harvest rates on 5, 10, or 20-year intervals and maintains these rates until the next management intervention.

**Table 2.** Harvest control rules used in the management scenarios.*

| Harvest control rule (HCR) |
| --- |
| *Current fishing mortality*<br>This HCR continues the initial fishing mortality rate (i.e., $F$ in 2012) through all years. |
| *Economically optimal fishing mortality*<br>This HCR achieves maximum net present value (NPV) over an infinite time horizon under the current climate and biological conditions. Each stock has its own optimized harvest policy where fishing mortality rate is a function of biomass. This HCR is determined using a dynamic optimization routine for each stock. |
| *Gradual shift from current to open access fishing mortality*<br>This HCR is only relevant to transboundary stocks. For these stocks, fishing mortality begins at the initial fishing mortality rate (i.e., $F$ in 2012), then changes at a constant rate towards open access fishing mortality (i.e., fishing mortality that achieves open access equilibrium at 30% of $B_{MSY}$), which is reached in the year in which the first spatial shift into or completely out of an EEZ occurs. Fishing mortality remains at the open access rate for all subsequent years. |
| *Gradual shift from economically optimal to open access fishing mortality*<br>This HCR is only relevant to transboundary stocks. For these stocks, fishing mortality begins at the economically optimal level given biomass in 2012, then changes at a constant rate towards open access fishing mortality (i.e., fishing mortality that achieves open access equilibrium at 30% of $B_{MSY}$), which is reached in the year in which the first spatial shift into or completely out of an EEZ occurs. Fishing mortality remains at the open access rate for all subsequent years. |

\* See the Gaines et al. [19] supplementary information for more details on the management scenarios and harvest control rules.

### Country-level fisheries outcomes

We evaluated the impact of climate change and management reform on the fisheries of 156 coastal sovereign countries summing across their domestic and territorial exclusive economic zones (EEZs). We scaled the projections of Gaines et al. [19] from the global- to country-level by assuming that the proportion of a species’ overall range occurring inside a country’s EEZ is identical to the proportion of the species’ overall carrying capacity occurring inside the country’s EEZ. This proportion was used to generate time series of biomass, harvest, and profit for each species in each country under all three emissions scenarios and five management scenarios. We summarized country-level projections by comparing fisheries outcomes: (1) in 2100 relative to today under each management scenario and (2) over the entire period (2012-2100) for each of the adaptation scenarios relative to the business-as-usual scenario. These approaches allow us to, respectively, estimate the projected impact of climate change on national fisheries outcomes under the different management scenarios and the cost of failing to adapt national fisheries management to account for climate change.

For Approach 1, we compared the percent difference in harvests and profits in 2100 relative to today (i.e., 2012) under each management scenario. While Gaines et al. [19] performed this comparison using only the projection endpoints (i.e., values in 2012 and 2100), we compared mean decadal values at the ends of the projection window (i.e., mean value in 2012-2021 and 2091-2100) to reduce sensitivity to specific endpoint values. For Approach 2, we compared the percent difference in cumulative harvest and cumulative profits between the four adaptation scenarios and the business-as-usual scenario. By examining differences in cumulative harvest and profits, this approach is also insensitive to endpoints and documents the accumulated benefits or losses of climate adaptive management. In both approaches, we quantified the impact of climate change and fisheries management on fisheries health as the mean proportion of stocks with biomass above B_MSY_, the biomass that produces MSY when fished at F_MSY_, by century’s end (2091-2100). This is a common target for fisheries management (i.e., U.S. Magnuson Stevens Act, E.U. Common Fisheries Policy, and U.N. Sustainable Development Goals). This performance metric better reflects the goals of fisheries management than percent change in biomass. For example, decreasing biomass in a previously undeveloped fishery is an expected consequence of economically optimal management and should only be perceived negatively when the decrease reduces biomass below the target.

## Results

### Impacts of climate change on maximum sustainable yield

Maximum sustainable yield (MSY) of the evaluated stocks is forecast to decrease by 2.0%, 5.0%, and 18.5% from 2012-2021 to 2091-2100 under RCPs 4.5, 6.0, and 8.5, respectively (Figure 1). Note that these values differ slightly from those reported in Gaines et al. [19] because we excluded mixed-species stocks and measured changes in MSY using decadal means. Across emissions scenarios, MSY is projected to decrease for equatorial countries and increase for poleward countries (Figure 1). Particularly dramatic reductions in MSY are predicted for the equatorial West African countries. Even under the least severe emissions scenario, nineteen countries, fifteen of which are in West Africa, are projected to experience reductions in MSY of 50-100%. The number of countries projected to experience dramatic losses in MSY, and the intensity of these losses, expands under the more severe emissions scenarios. In the most severe scenario, 51 countries are expected to experience reductions in MSY of 50-100% (Figure 1). All eighteen West African countries south of Senegal and north of Angola (including these two countries) are forecast to experience reductions in MSY greater than 85%. The equatorial Indo-Pacific and South America are also projected to experience considerable losses in MSY under the three emissions scenarios, with especially pronounced losses under RCP 8.5 (Figure 1). Twenty-two countries are projected to experience increases in MSY under all three emissions scenarios with seven of these countries showing a 15% average increase in MSY across scenarios. The five most consistent and pronounced climate change “winners” are: Finland, Antarctica, Norway (4 EEZs: Norway plus Bouvet Island, Jan Mayen, and Svalbard), Portugal (3 EEZs: Portugal plus Azores and Madeira), and Fiji.

**Figure 1.**
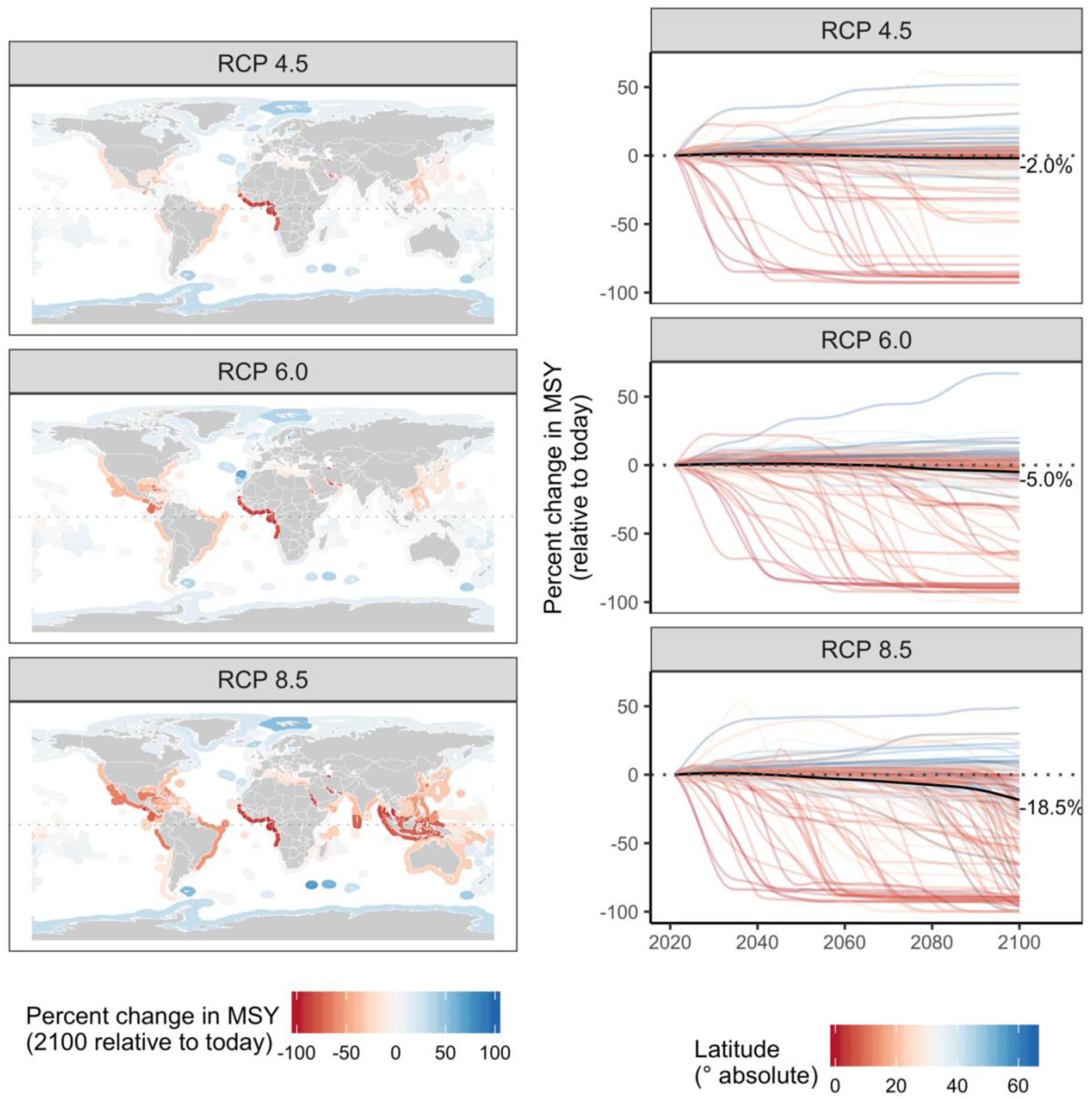
Percent change in maximum sustainable yield (MSY) under each emission scenario. In the left column, maps show the percent change in MSY from 2012-2021 (“today”) to 2091-2100 in each exclusive economic zone. In the right column, the colored lines show the percent change in MSY (measured in 10-year running averages) relative to 2012-2021 (“today”) for each of 156 countries and the black lines show the percent change globally.

### Ability for management reform to mitigate global climate impacts

Business-as-usual (BAU) management results in both lower catches and profits in the future relative to today under all three emissions scenarios (Figure 2). In contrast, full adaptation yields both higher catches and profits in the future in all but the most severe emissions scenario (RCP 8.5); in this scenario, full adaptation yields higher profits but lower catches in the future relative to today. Addressing productivity shifts and range shifts in isolation is insufficient for jointly maintaining catch and profits into the future under any of the emissions scenario (Figure 2). However, realistic adaptation, which recalibrates productivity management at 5, 10, and 20-year intervals and maintains this management regime as stocks shift across boundaries, frequently achieves better outcomes in the future relative to today (Figure 2). Notably, realistic adaption that implements adaptive management at 5-year intervals performs comparably to full adaptation and generates both higher catch and profits in the future relative to today under the two least severe emissions scenarios (Figure 2).

**Figure 2.**
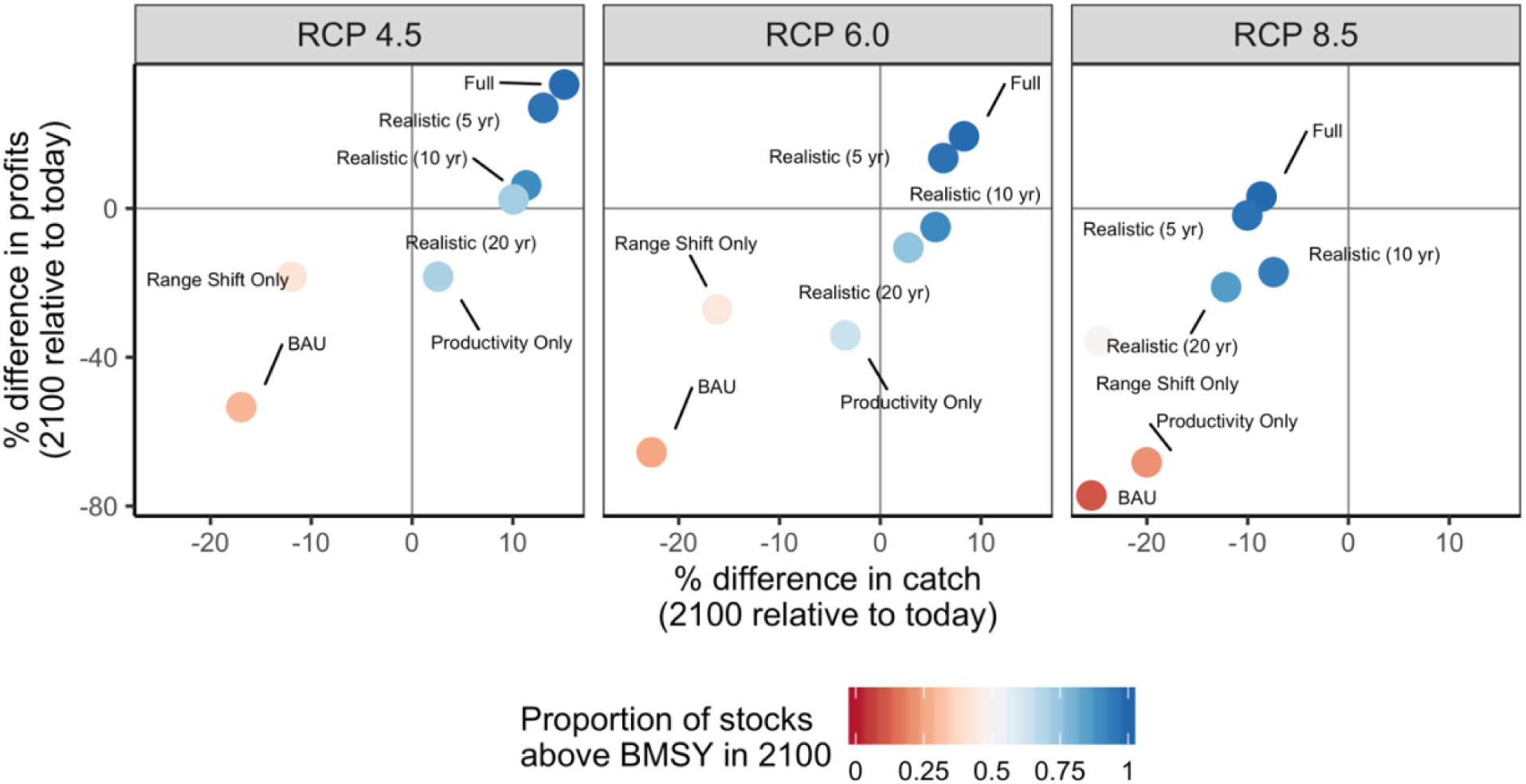
Percent difference in mean catch and profits in 2091-2100 relative to 2012-2021 (“today”) from all stocks under each emission and management scenario.

### Ability for management reform to mitigate country-level climate impacts

While business-as-usual management results in lower catches and profits relative to today for the majority of countries (82-85% of countries), full adaptation yields higher catches and profits for a majority of countries in all but the most severe emission scenario (Figures 3 and S1). In this scenario, only 35% of countries experience both increased profits and catches, while 59% of countries experience both reduced catches and profits (Figure 3). Realistic adaptation implemented at 5-year intervals achieves outcomes quite similar to full adaptation: it results in higher catch and profits for the majority (56-63%) of countries under RCPs 4.5 and 6.0 but lower catch and profits for the majority (59%) of countries under RCP 8.5 (Figure 3). The ability for adaptation to maintain or increase fisheries outcomes under climate change is sensitive to the direction and magnitude of changes in underlying productivity (Figures 3-5). For example, the West African countries projected to experience the greatest losses in MSY are also projected to have the most limited ability to mitigate these impacts (Figures 1 and 4). Although realistic adaptation (5-yr) could increase both catch and profits for 51% of the countries projected to lose underlying productivity (i.e., lower MSY) in the least severe emissions scenario, it could increase outcomes despite losses in productivity for only 23% of countries in the most severe emissions scenario (Figure 4). In comparison, realistic adaptation (5-yr) could increase both catch and profit for a much larger proportion of countries projected to gain underlying productivity: 78% of these countries (n=69) could increase both catch and profits in the least severe emissions scenario and this percentage actually increases to 95% in the most severe emissions scenario as these poleward countries (n=22) inherit even more productivity (Figure 4). Neither realistic (5-yr) nor full adaptation are sufficient to maintain fisheries outcomes into the future for all countries, but they are nearly always preferable to business-as-usual management. In all but the most severe emissions scenario, both full adaptation and realistic adaptation yield both higher cumulative catches and profits than business-as-usual management for nearly all countries (98-99% of countries; Figure 6). In the most severe scenario, full adaptation and realistic adaptation yield higher cumulative profits than business-as-usual management, but achieve lower cumulative catches for 40-41% of countries (Figure 6).

**Figure 3.**
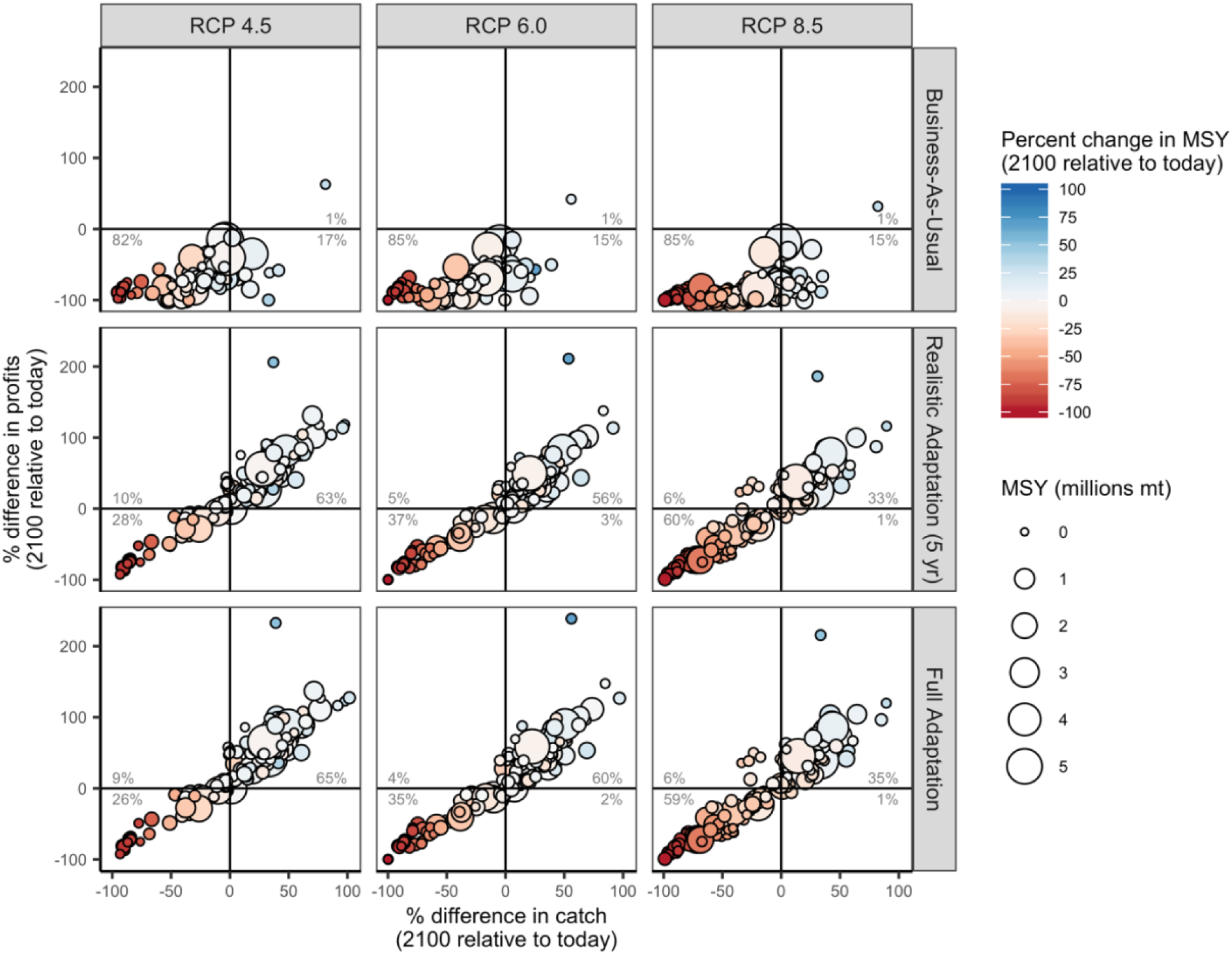
Percent difference in mean catch and profits in 2091-2100 relative to 2012-2021 (“today”) for 156 countries under three emissions scenarios (columns) and three management scenarios (rows). The percentage labels indicate the percentage of countries falling in each quadrant of catch and profit outcomes.

**Figure 4.**
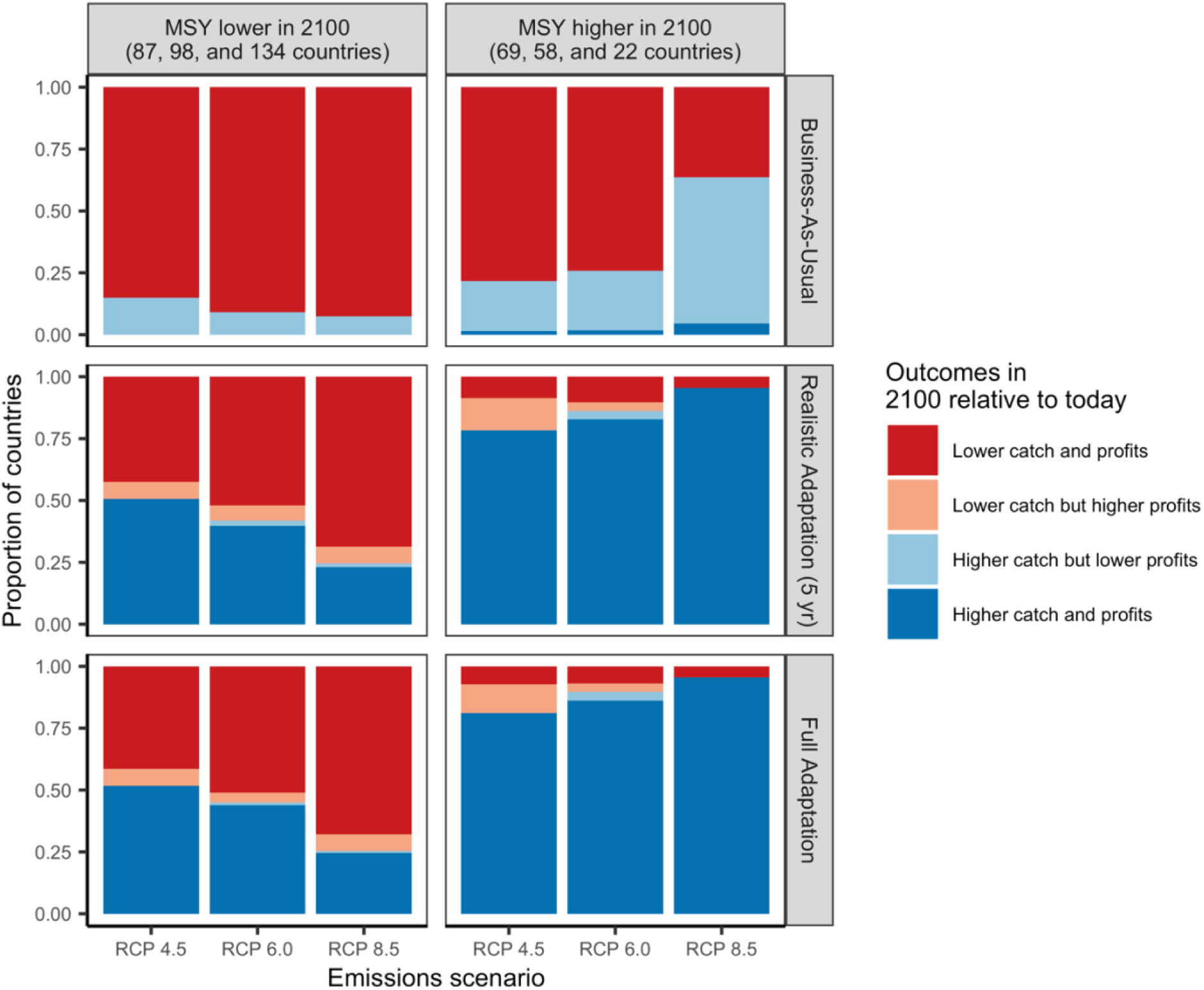
Influence of changes in maximum sustainable yield (MSY) on the ability for management to generate higher catch and profits in the future (2091-2100) relative to today (2012-2021). Bars indicate the proportion of countries experiencing each combination of catch and profits trajectories under each emissions scenario, management scenario (rows), and change in underlying productivity (columns). The number of countries experiencing reductions in MSY increases under increasingly severe emissions scenarios (see column title for numbers). Although the number of countries experiencing gains in MSY decreases under increasingly severe emissions scenarios (see column title for numbers), the gains in MSY in these countries are actually magnified with increasing emissions (i.e., more fish stocks move into their exclusive economic zones with more rapid warming).

**Figure 5.**
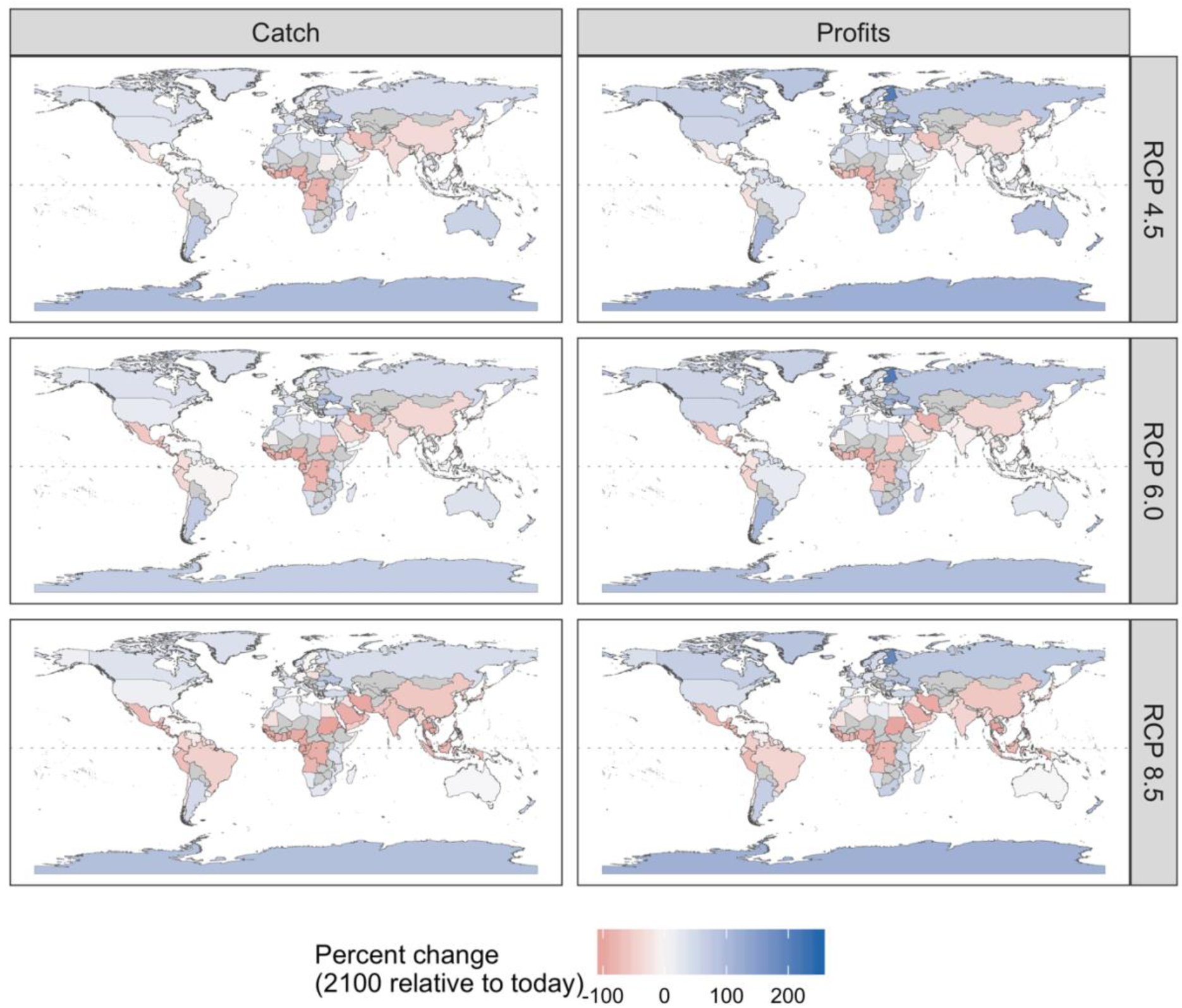
Percent difference in mean catch and profits in 2091-2100 relative to 2012-2021 (“today”) for 156 countries under realistic adaptation implementing management at 5-year intervals. Grey shading indicates countries without marine territories.

**Figure 6.**
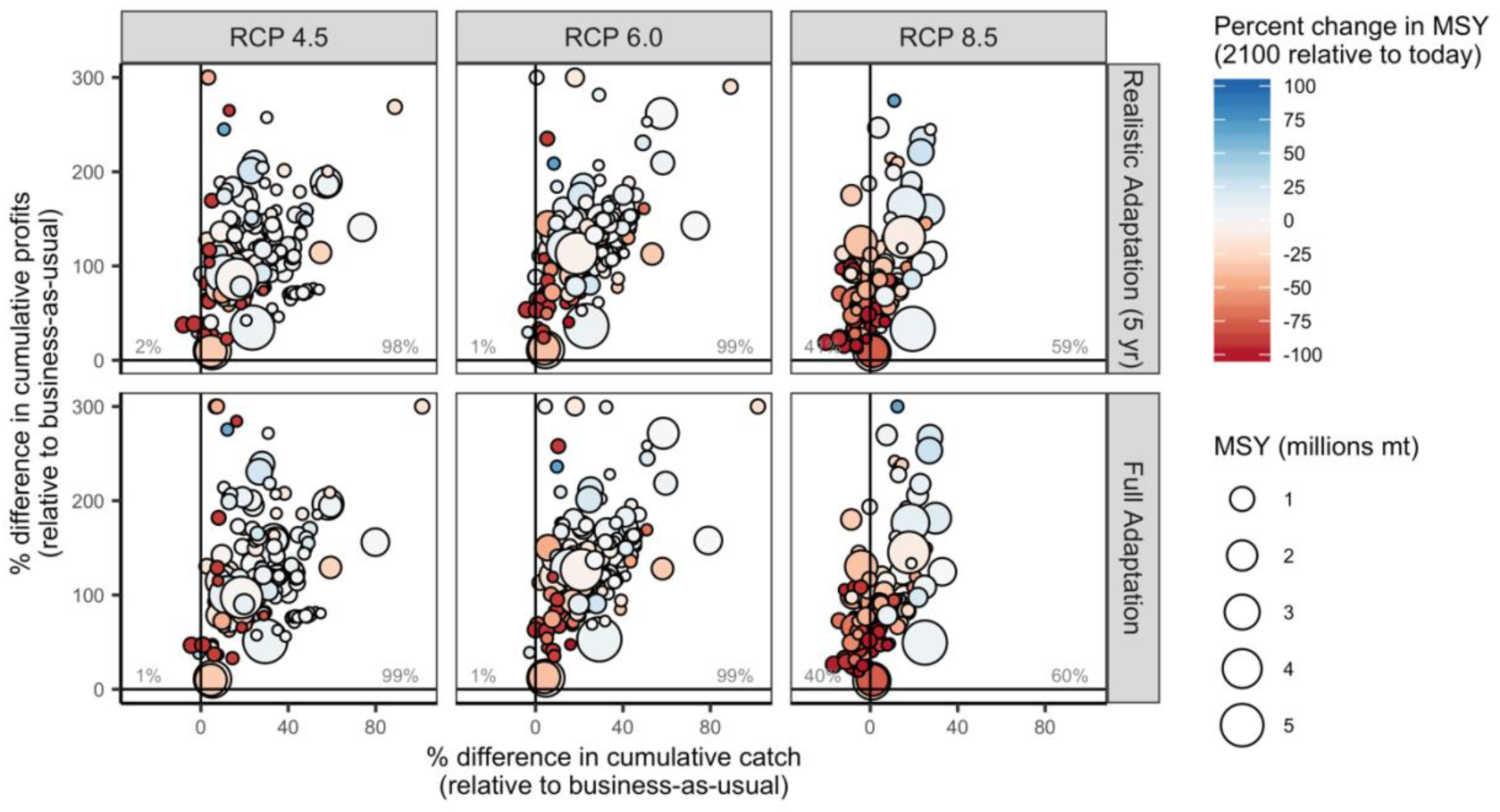
Percent difference in cumulative catch and cumulative profits from 2012-2100 relative to business-as-usual for 156 countries under three emissions scenarios (columns) and two adaptation scenarios (rows). The percentage labels indicate the percentage of countries falling in each quadrant of catch and profit outcomes.

## Discussion

Overall, our results indicate that climate change will dramatically alter the distribution and productivity of marine fisheries, but plausible climate-adaptive management reforms could minimize or eliminate negative impacts in most countries. This reinforces and expands upon the work of Gaines et al. [19] in two important ways. First, whereas Gaines et al. [19] document the benefits of management reform at a global-level, we focus on the distributional consequences of these global effects by evaluating the benefits that individual countries stand to gain from climate-adaptive fisheries reforms. Second, we recognize that perfectly adapting to changing productivity will be a challenge in even the most sophisticated fisheries systems [15, 28] and evaluate a more realistic scenario that implements well-intentioned, yet imperfect, adaptation to productivity shifts. These expansions are important because they place more realistic bounds on the ability for management to mitigate the impacts of climate change and present practitioners with a tool for investigating the impacts of climate change and opportunities for reform in their respective country’s fisheries.

Our model predicts shifts in productivity that are consistent in both pattern and magnitude with a recent ensemble model [5] that averages the predictions of six other peer-reviewed marine ecosystem models. We estimated 2.0% and 18.5% decreases in maximum sustainable yield from 2012-2100 under RCPs 4.5 and 8.5, respectively. By comparison, Lotze et al. [5] estimated 8.6% (±6.0% SD) and 17.2% (±10.7% SD) decreases in marine animal biomass in the absence of fishing from 1990-2100 under the same two emissions scenarios. The Lotze et al. [5] ensemble model, its constituent models, and our model all predict increases in productivity in poleward regions and decreases in productivity in tropical to temperate regions. The slight differences in the productivity shifts predicted by our model and the ensemble model are unsurprising given the differences in the structure, mechanistic drivers, and taxonomic scope of our model and the ensemble’s constituent models.

Importantly, however, our approach differs from these studies, because, in addition to forecasting the impact of climate change on the biological potential of fisheries, we consider the impact of alternative human responses to these changes, which could either exacerbate or alleviate the impacts of changing biological potential [13]. Indeed, our results indicate that all countries would benefit from reforming current management to account for shifting distributions and productivity and that many countries could even see higher catch and profits than today with such reforms. However, the ability for management reform to mitigate the impacts of climate change is dependent on swift efforts to reduce greenhouse gas emissions. Even perfect climate-adaptive management (“full adaptation”) is unable to maintain current catch and profits under high-end greenhouse gas emissions (RCP 8.5). Furthermore, although perfect adaptation could maintain global catch and profits under partial emission reductions (RCP 6.0), tropical and temperate regions would still incur dramatic losses in fisheries benefits. This underscores the fact that even emission reductions consistent with the Paris Agreement could have significant impacts on the ability for fisheries to feed and employ people into the future [29, 30].

The development and implementation of stock assessment methods and management strategies necessary to achieve benefits even in the face of climate change is nascent but rapidly developing. For example, Skern-Mauritzen et al. [31] reviewed 1,250 stock assessments from around the world and found that only 2% incorporated ecosystem information into tactical management advice. In the United States, Marshall et al. [21] used a broader definition of inclusion and ecosystem information and found greater, though still limited, incorporation of ecosystem information into stock assessments: 24% of 206 evaluated assessments included ecosystem information in the assessment report. The effective incorporation of environmental information into management strategies is similarly challenging but is also increasing in frequency and effectiveness. Punt et al. [28] reviewed management strategy evaluation (MSE) studies that test procedures for setting environmentally-linked harvest control rules and found that, in general, these procedures were only effective when the environmental drivers were well understood. This emphasizes the need for increasing monitoring and process-oriented lab and field studies in conjunction with the development and testing of more sophisticated analytical techniques [32].

Furthermore, achieving the benefits of climate-adaptive fisheries reform will require accounting for shifting productivity and distributions along a gradient of scientific, management, and enforcement capacities. Many countries lack the monitoring programs required to detect and describe shifts in distribution and productivity, the scientific capacity for conducting either climate-agnostic or climate-adaptive stock assessments, and the management capacity for setting and enforcing fisheries regulations [25,33,34]. This is frequently the case for the tropical developing countries that are forecast to experience the greatest losses in fisheries catch and profits under climate change and exhibit the greatest vulnerability to these reductions in food and income [35]. The tools for enacting climate-adaptive fisheries reforms and achieving biological and socioeconomic resilience to climate change will have to span this gradient of capacity.

Fortunately, a growing body of literature provides guidance on accounting for shifting distributions and productivity in fisheries assessment and management [14,17,36,37] and for fostering socioeconomic resilience to climate change [38–40] in diverse fisheries systems. In the remainder of this paper, we provide a brief overview of this literature and recommend general principles as well as specific strategies for achieving the benefits of climate-adaptive management reforms. We offer recommendations for higher and lower capacity fisheries systems as well as recommendations for countries where even the best management reforms will be unable to offset the negative impacts of climate change.

### Guiding principles for climate-adaptive fisheries management

#### Principle #1: Implement best practices in fisheries management

Historically, well-managed fisheries have been among the most resilient to climate change [4], and our results predict that well-intended, albeit imperfect, management will continue to confer climate resilience. Together, these results indicate that the wider implementation of best practices in fisheries management will mitigate many of the negative impacts of climate change. In higher capacity systems, best practices include scientifically-informed catch limits, accountability measures, regional flexibility in policy practices, and protection of essential fish habitat [41]. In the United States, such measures have contributed to dramatic declines in overfishing, increases in biomass, and maintenance of catch and profits [42]. In lower capacity systems, best practices include implementing “primary fisheries management” [43] that uses best available science and precautionary principles to manage data-poor and capacity-limited fisheries and establishing local, rights-based management [39] to incentivize sustainable stewardship. Rights-based management systems include catch share programs such as Individual Transferable Quotas (ITQs) and Territorial Use Rights in Fisheries (TURFs) that define property rights over catch and space, respectively [44]. By giving users ownership of the resource, well-designed rights-based management systems incentivize long-term stewardship and have been shown to promote compliance, prevent overfishing, and increase profits [16,45,46]. Enforcement and the strength of fishing pressure limits are also key for successful fisheries management [47] and contribute to a precautionary approach in the face of climate change. Overall, fisheries best practices confer ecological resilience by maintaining healthy stock sizes, age structures, and genetic diversity and socioeconomic resilience by providing a portfolio of options to fishers and a buffer against climate-driven losses in any one target stock.

#### Principle #2: Be dynamic, flexible, and forward-looking

Adapting to climate change will require dynamic, flexible, and forward-looking management. This can be achieved by aligning management policies with the spatio-temporal scales of climate change, ecosystem change, and socioeconomic responses [14]. In higher capacity systems, this could involve four broad strategies. First, managers can envision and prepare for alternative futures using tools such as forecasts [48], structured scenario planning [49], holistic ecosystem models [50], risk assessments [51], and climate vulnerability analyses [52]. Second, the proliferation of near real-time biological, oceanographic, social, and/or economic data can be harnessed for proactive and dynamic adjustments in spatial and temporal management actions [53]. Third, developing harvest control rules that account for or are robust to changing environmental conditions affecting productivity can increase catch while also reducing the probability of overfishing [54]. Finally, all of these management procedures should be simulation tested through management strategy evaluation (MSE; [55]) to measure the efficacy of alternative strategies and their robustness under different climate scenarios [28]. In lower capacity systems, forward looking fisheries management could include precautionary management to buffer against uncertainty [56] as well as management strategies that preserve population resilience, age structure, and genetic diversity. For example, size limits, seasonal closures, and protected areas can be used to protect the big, old, fecund, females (BOFFs) that disproportionately contribute to reproductive output [57] and to maintain the genetic diversity required to promote evolutionary adaptations to climate change.

#### Principle #3: Foster international cooperation

Shifting distributions are already generating management challenges and the rates of these shifts and associated conflicts are expected to increase with climate change [17,18,58]. New or strengthened international institutions and agreements will be necessary to ensure that management remains sustainable as stocks shift between jurisdictions. First, this will require sharing data between Regional Fisheries Management Organizations (RFMOs) or countries to identify, describe, and forecast shifting stocks. Second, it will require a commitment to use these shared data to inform collaborative management. For example, these data could be used to regularly and objectively update national allocations of catch or effort based on changes in distribution rather than historical allocations (e.g., [59, 60]). An alternative approach could be to develop fisheries permits that are tradeable across political boundaries, which would allow future resource users access to fisheries not yet in their waters and incentivize good management [61]. Finally, incentivizing the cooperation necessary to establish data sharing and collaborative management will require overcoming prevailing management mentalities that one party “wins” while the other “loses” when stocks shift across boundaries. This could involve broadening negotiations to allow for alternative avenues of compensation or “side payments” [62]. In cases where establishing international cooperation proves difficult, marine protected areas (MPAs) placed along country borders could buy time for negotiations by protecting stocks as they shift across borders. A more precautionary approach would be to put new fishing areas on hold until adaptive management can be put in place, as illustrated by the Central Arctic Ocean Fisheries Agreement (e.g., the CAOF Agreement, [63]).

#### Principle #4: Build socioeconomic resilience

The impact of climate change on fishing communities can be reduced through measures that increase socioeconomic resilience and adaptive capacity to environmental variability and changing fisheries [40,64,65]. Across low to high capacity systems, these measures include (1) policies that facilitate flexibility, such as diversification of access to fisheries and alternative livelihoods, (2) policies that provide better assets, such as the enhancement of fisheries technology and capacity, (3) policies that provide better organization in the system, including multi-level governance, community-based management, and other governance structures [14, 39], and (4) policies that promote agency and learning [40]. For example, policies that promote access to multiple fisheries provide fishers with a portfolio of fishing opportunities that can buffer against variability [66, 67] while policies that promote diverse livelihoods reduce reliance on fisheries [68, 69]. Increased mobility through technological enhancements can increase social resilience by allowing fishers to follow shifting stocks [40], but can also result in the migration of fishers. Multi-level governance promotes flexibility in resource governance by matching biological and management across scales [70]. Community-based management can increase adaptive capacity by incorporating local knowledge and can improve sustainability by fostering a sense of stewardship [71]. Spatial-rights based approaches such as TURFs may confer social resilience insofar as they are often community-managed and allow fishers to generate revenues through other compatible activities such as tourism, recreation, and aquaculture [72]. On the other hand, ITQs may confer a different kind of resilience because rights are defined over fish catch, not spatial areas, so they may be more resilient to range shifts arising from climate change. Furthermore, all of these measures can be designed to reduce fishing pressure, and promote ecological resilience to climate change.

### Aquaculture could help compensate for losses in capture fisheries

Even the best climate-adaptive management will be unable to maintain current catch and profits in most tropical developing countries. Although these countries should still pursue climate-adaptive reforms to maximize catch and profits from capture fisheries, they will also need to develop, expand, and reform other sectors to compensate for capture fishery losses and meet growing production demands [73]. Marine aquaculture (hereafter called mariculture), the cultivation of marine animals and plants, presents a particularly promising substitute for capture fisheries. The biological potential for mariculture is enormous [74] and exceeds both current production and projected demand even after accounting for economic feasibility and the availability of feed for fed-finfish mariculture [75]. This potential is expected to decrease under climate change [76] but breeding a larger proportion of stocks for fast growth could more than offset these negative impacts [77]. Although mariculture has the potential to feed millions of people, it also poses a number of environmental problems including pollution, habitat conversion, disease and parasite transmission, and escapement and hybridization [78]. The expansion of large-scale mariculture for increased food and employment opportunities will thus require a better understanding of these environmental tradeoffs and the best practices for managing them [79].

## Conclusions

Although climate change is expected to reduce the productivity of marine fisheries globally [5], climate-adaptive fisheries management reforms could mitigate many of the negative impacts on the food and income provisioning potential of the ocean [19]. Our results suggest that climate-adaptive fisheries could result in higher catch and profits than business-as-usual management in all countries. For most countries, climate-adaptive management reforms could result in higher catch and profits in the future than today. However, the ability for management reforms to offset negative impacts is diminished under increasingly severe greenhouse gas emission scenarios. Thus, swift actions to reduce emissions will be necessary to limit the impacts of climate change on fisheries, especially in developing tropical countries. For many of these countries, even the best climate-adaptive fisheries reforms will be insufficient to maintain current levels of catch and profits into the future. Adaptation in these countries will require innovations in sustainable mariculture and other food sectors to ensure that countries are able to meet the food and nutrition requirements of their growing populations [73]. As land-based sources of food also falter [80], the ocean will become an increasingly important source of nutrition. Achieving these benefits will depend on swift and innovative management actions.

## Acknowledgements

This work was funded by the Environmental Defense Fund. J.G.M. is supported by the “Tenure-Track System Promotion Program” of the Japanese Ministry of Education, Culture, Sports, Science and Technology (MEXT). E.O. is funded by the European Research Council project CLOCK (GA. 679812) and GAIN-Xunta de Galicia Oportunius program. We thank Merrick Burden, Kate Bonzon, and Willow Battista for valuable discussions while preparing the manuscript.

---

**Figure S1.**
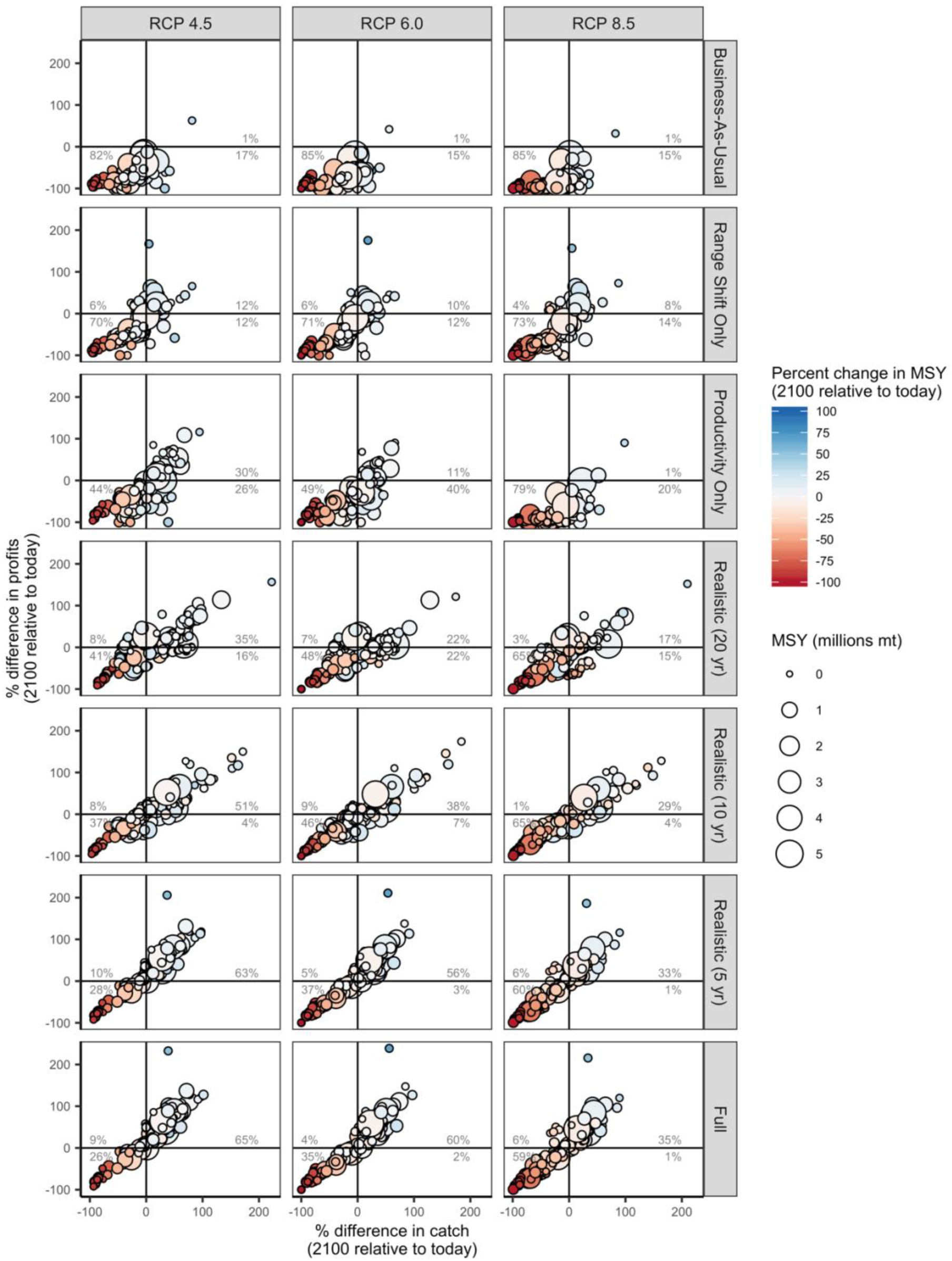
Percent difference in mean catch and profits in 2091-2100 relative to 2012-2021 (“today”) for 156 countries under three emissions scenarios (columns) and seven management scenarios (rows). The percentage labels indicate the percentage of countries falling in each quadrant of catch and profit outcomes.

**Figure S2.**
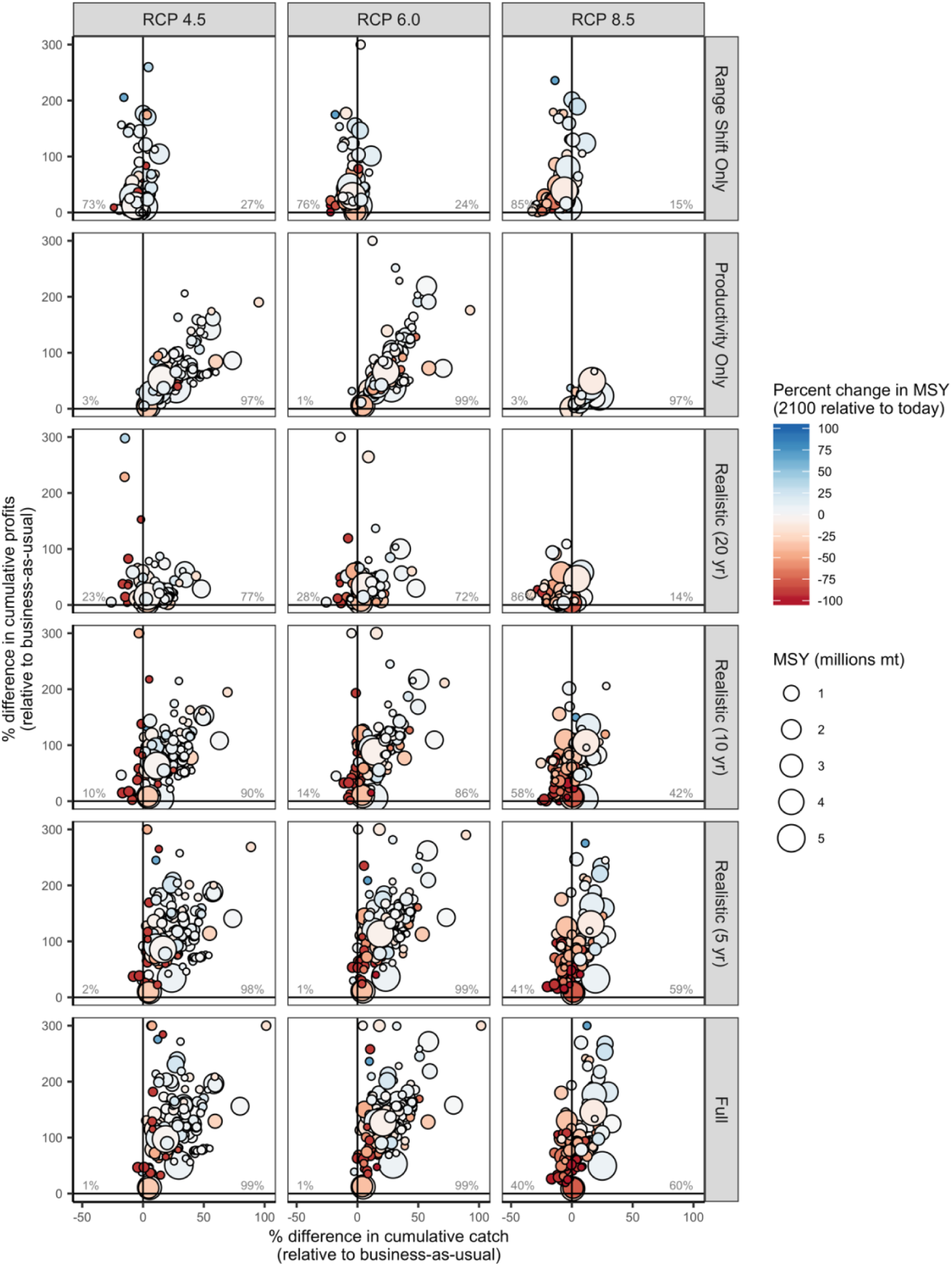
Percent difference in cumulative catch and cumulative profits from 2012-2100 relative to business-as-usual for 156 countries under three emissions scenarios (columns) and six management scenarios (rows). The percentage labels indicate the percentage of countries falling in each quadrant of catch and profit outcomes.

**Table S1.**
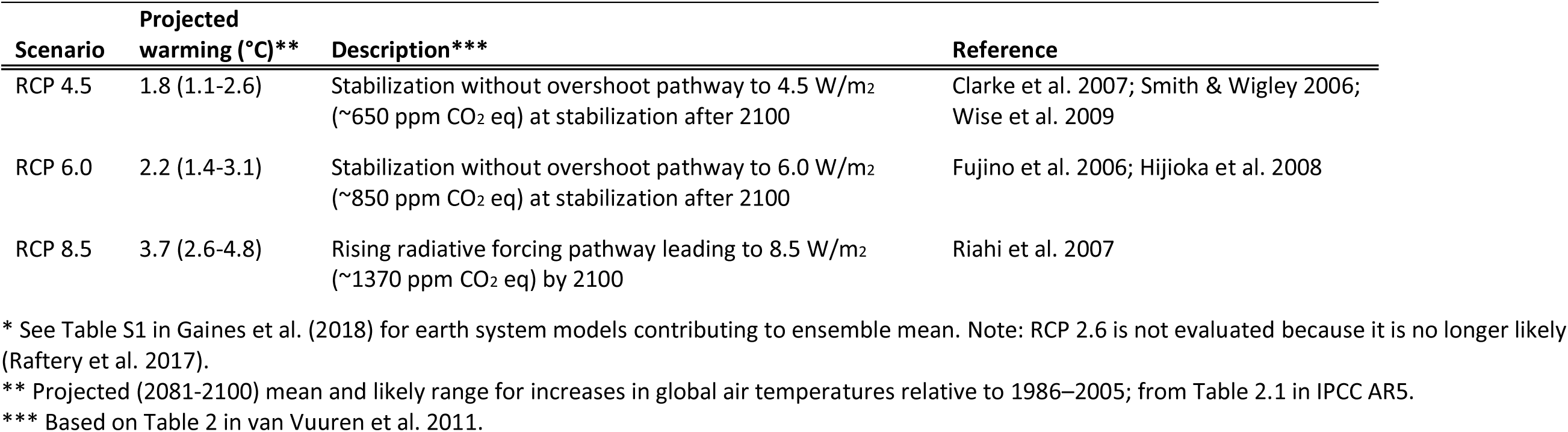
Representative Concentration Pathways (RCPs) evaluated in the analysis*.

